# Pulmonary ductal coarctation and left pulmonary artery interruption; pathology and role of neural crest and second heart field during development

**DOI:** 10.1101/2020.01.17.910224

**Authors:** Adriana C. Gittenberger-de Groot, Joshua C. Peterson, Lambertus J. Wisse, Arno A.W. Roest, Robert E. Poelmann, Regina Bökenkamp, Nynke J. Elzenga, Mark Hazekamp, Margot M. Bartelings, Monique R.M. Jongbloed, Marco C. DeRuiter

**Affiliations:** Department of Cardiology, Leiden University Medical Center, Leiden, the Netherlands; Department of Anatomy & Embryology, Leiden University Medical Center, Leiden, the Netherlands; Department of Pediatric Cardiology, Leiden University Medical Center, Leiden, the Netherlands; Department of Animal Sciences and Health, Leiden University, Leiden, the Netherlands; Department of Pediatric Cardiology, University Hospital Groningen, Groningen, the Netherlands; Department of Cardio-thoracic Surgery, Leiden University Medical Center, Leiden, the Netherlands

**Author notes:** Corresponding author: Adriana C. Gittenberger-de Groot.

**Keywords:** cardiac malformations, cardiac development, ductus arteriosus, pulmonary atresia, left pulmonary artery, tetralogy of Fallot, 22q11 deletion syndrome, juxtaductal pulmonary coarctation

## Abstract

**Objectives:** In congenital heart malformations with pulmonary stenosis to atresia an abnormal lateral ductus arteriosus to left pulmonary artery connection can lead to a localised narrowing (pulmonary ductal coarctation) or even interruption We investigated embryonic remodelling and pathogenesis of this area.

Material and methods. Normal development was studied in WntCre reporter mice (E10.0-12.5) for neural crest cells and Nkx2.5 immunostaining for second heart field cells. Data were compared to stage matched human embryos and a VEGF120/120 mutant mouse strain developing pulmonary atresia.

**Results:** Normal mouse and human embryos showed that the mid-pharyngeal endothelial plexus, connected side-ways to the 6^th^ pharyngeal arch artery. The ventral segment formed the proximal pulmonary artery. The dorsal segment (future DA) was solely surrounded by neural crest cells. The ventral segment had a dual outer lining with neural crest and second heart field cells, while the distal pulmonary artery was covered by none of these cells. The asymmetric contribution of second heart field to the future pulmonary trunk on the left side of the aortic sac (so-called pulmonary push) was evident. The ventral segment became incorporated into the pulmonary trunk leading to a separate connection of the left and right pulmonary arteries. The VEGF120/120 embryos showed a stunted pulmonary push and a variety of vascular anomalies.

**Summary:** Side-way connection of the DA to the left pulmonary artery is a congenital anomaly. The primary problem is a stunted development of the pulmonary push leading to pulmonary stenosis/atresia and a subsequent lack of proper incorporation of the ventral segment into the aortic sac. Clinically, the aberrant smooth muscle tissue of the ductus arteriosus should be addressed to prohibit development of severe pulmonary ductal coarctation or even interruption of the left pulmonary artery.

## Introduction

Before birth the ductus arteriosus (DA) connects the pulmonary trunk (PT) to the descending aorta, thus bringing oxygen-rich placental blood to the systemic circulation, largely circumventing the still not functioning lungs. In the perinatal period the muscular DA starts physiological closure by contraction followed by anatomical sealing and subsequent ligament formation [1]. Some heart malformations present with narrowing of the pulmonary outflow tract (OFT) as in tetralogy of Fallot with severe pulmonary stenosis or pulmonary atresia and a ventricular septal defect (VSD). Here, instead of being in direct continuity with the PT (Figure 1a), we can find an abnormal lateral DA to left pulmonary artery connection [2, 3]. Often a narrowing in the proximal part of the left pulmonary artery, a so-called pulmonary ductal coarctation (PDC), is seen (Figure 1b). It is not known whether this leftward positioning of the DA connection is a developmental anomaly or whether it reflects the persistence of a normal embryonic stage as seen in an evo-devo setting in chicken embryos [4, 5, 6].

**Figure 1:**
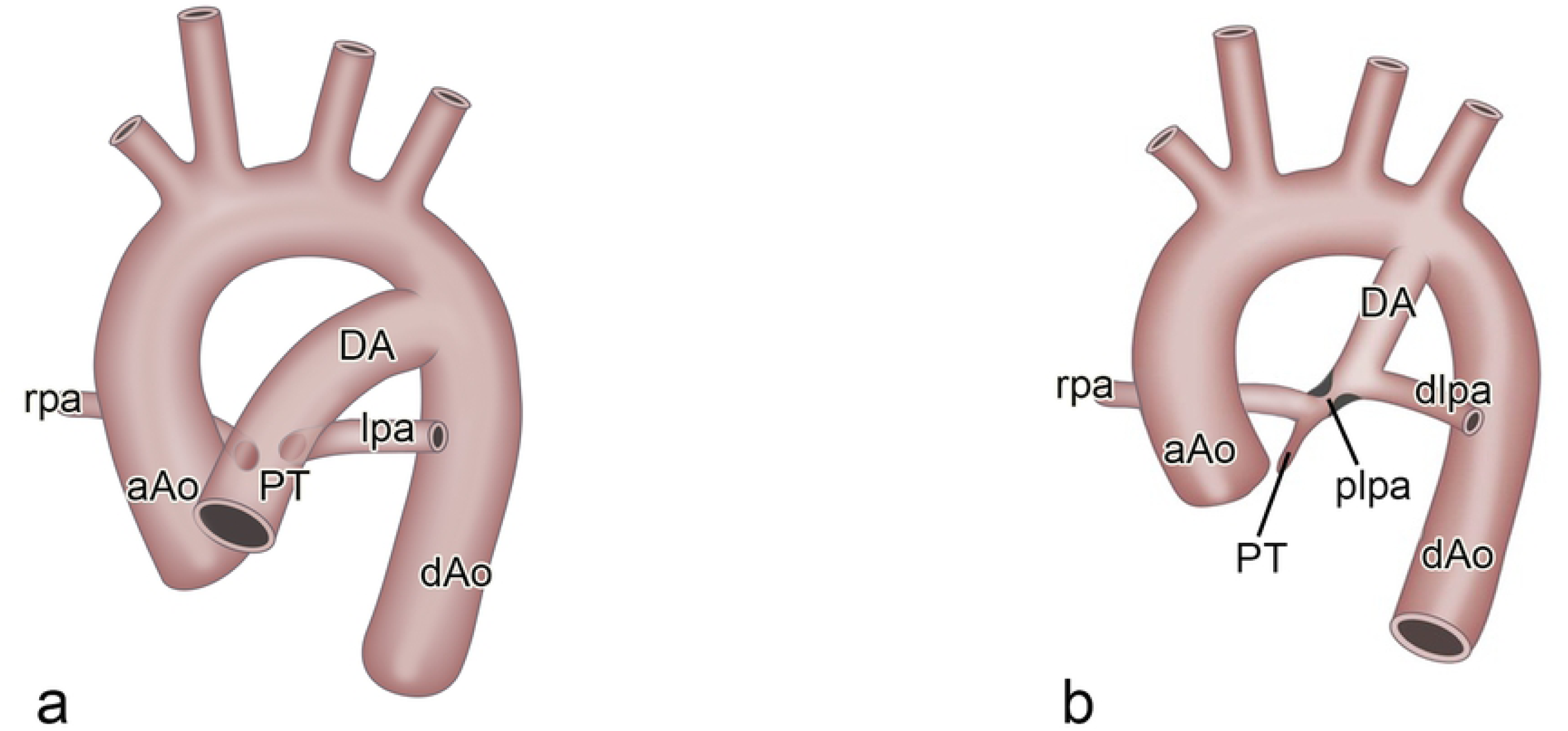
**a. S**chematic view of the normal perinatal aortic arch showing the ascending aorta (aAo) and the pulmonary trunk (PT). De ductus arteriosus (DA) connects the PT with the descending aorta (dAo). The left (lpa) and right (rpa) pulmonary arteries are dorsally hooked up to the PT. Fig **b**. Schematic representation of a case with an abnormal lateral DA to lpa connection creating a proximal (plpa) and distal (dlpa) part. Aberrant DA tissue (black indentations), extending into the plpa, create the narrowing hallmark of the pulmonary ductal coarctation.

Formation of the pharyngeal arch artery (PAA) system, with merging of ventral and dorsal sprouts to form a complete arch, has been extensively studied in the human embryo [7]. Refinements were brought by acknowledging the special status of the 6^th^ or pulmonary arch artery [8] in which it was shown that the anlage of the pulmonary arteries, derived from the endothelium of the mid-pharyngeal endothelial strand (MPES), does not grow out towards the lung but connects to the ventral sprout before the completion of the 6^th^ PAA. On the left side the dorsal segment of the 6^th^ PAA persists as the DA while this segment on the right side disappears [7] by a process of selective apoptosis [9]. To achieve the perinatal situation in which the DA connects directly to the pulmonary trunk (Figure1a) separately from the adjoining left and right pulmonary arteries, the ventral sprout of the 6^th^ PAA either has to become part of the pulmonary artery or has to disappear. The remodelling in this area has not been adequately studied but is relevant for understanding the abnormal connection of the DA to the left pulmonary artery as encountered in PDC (Figure 1b).

The normal remodelling of the PAAs takes place after the endothelium-lined vascular network is consolidated by smooth muscle cells [9, 10] derived from the surrounding mesoderm (second heart field / SHF) and mesectoderm (neural crest cells / NCCs). In order to better understand both normal and abnormal development in this complicated area we have applied in this study the more recent approach of investigating these cellular constituents.

Animal models, initially with the chimeric and retroviral tracing techniques in avian embryos and more recently transgenic reporter studies in mouse [11, 12, 13, 14] contribute to our understanding of the importance of the NCC population in PAA formation and OFT septation. Detailed information on the differential contribution of NCC to the wall of the ascending aorta and pulmonary trunk is emerging [14]. A relatively novel cell population, that received a hitherto neglected role, is the SHF [15]. This population is actually instrumental in adding myocardium to the right side of the heart [16] but it has also been described to contribute to the smooth muscle cells of the aortic and pulmonary wall [17]. We have previously shown that the contribution of the SHF is more prominent to the pulmonary side of the OFT, a process dubbed as the pulmonary push [18], being important for OFT septation and positional shift (often referred to as rotation) of the great vessels. It is of particular interest that this left-sided SHF population encircles the region of the left pulmonary artery-DA connection in which also NCCs are abundantly present.

Hardly any attention has been given to the specific contribution of the various cell types to vascular wall formation on boundaries at the merging of the 6^th^ PAA to the pulmonary system.

The abnormal lateral DA to left pulmonary artery connection is clinically relevant. In patients with pulmonary atresia and an intracardiac shunt, including cases with tetralogy of Fallot and univentricular hearts, DA closure potentially causes proximal left pulmonary artery stenosis or even interruption of the left pulmonary artery [2, 19, 20]. These additional anomalies complicate the treatment course of the patients seriously. Often it is necessary to address the stenotic proximal left pulmonary artery surgically [3] or by repeated catheter intervention because of the somatic growth of the patient.

In the present study a combined approach has been adopted to study this problem using both staged mouse embryos (normal, reporter mice and growth factor deficient mice) and stage-matched human embryos. Histopathology and clinical images of human patients will be presented to underscore the clinical importance.

## Materials and methods

### Embryonic material

The material used to study the development of the pulmonary arterial system consists of mouse embryos of various strains and stages, as well as human embryos from different sources. Reporter mice and immunohistochemistry have been used to study cellular constituents (NCCs, SHF mesoderm and endothelium) in arterial development. Growth factor deficient mice were studied for abnormal development. All mice were handled according to the Guide for Care and Use of Laboratory Animals (NIH), the experiments were approved by the LUMC Animal Welfare Committee (dec14184).

The following reporter mice (Black6 background using C57BL/6JLumc mice) derived from breeding strategies [14], have been used in this study: Wnt1Cre;mT/mG for NCC tracing; Mef2cCre;mT/mG for myocardial tracing (generous gift of Dr. Qingping Feng, The University of Western Ontario, London, Ontario, Canada). Furthermore, the VEGF 120/120 mouse, in which vascular endothelial growth factor (VEGF) isoforms 164 and 185 were blocked but an overexpression of soluble VEGF 120 is present [21, 22], was investigated for PAA pathological phenotypes [23]. Male and female mice were kept overnight and copulation was determined by the presence of a vaginal plug the following morning. Noon was considered as embryonic day (ED) 0.5. Embryos were removed at E10.0-E18.5 following dissection in phosphate buffered saline (PBS), pH 7.4.

Using upclimbing stages of reporter mice (E10.0-14.5) with Wnt/Cre for NCCs, PECAM for endothelium and Nkx2.5 staining for SHF-derived cells we re-investigated the connection of the 6^th^ PAA and the pulmonary arteries to the pulmonary trunk in detail. We compared normal mouse development with a series of normal human embryos from 4-24 mm crown-rump length (CR), which are overlapping with the studied mouse stages.

As a disease model we used the VEGF120/120 mutant mouse, originally presented as a model for the 22q11 deletion syndrome [24], with a spectrum of tetralogy of Fallot to pulmonary atresia [22]. It has already been described that in this mutant mouse initially the left and right 6^th^ PAA are formed and that, depending on the degree PT stenosis to atresia, the DA disappears during further embryonic development. This is accompanied in some cases with development of additional aorto-pulmonary connections [23].

Human embryos (4-24 mm CR length, Carnegie stages /CS12-24) were obtained as described [25, 26, 27], with the addition of one embryo from the Viennese department of Anatomy), fixed in 4% formaldehyde and serially sectioned.

### Postnatal material

From the human collection of neonatal heart malformations of the Leiden department of Anatomy, macroscopy and histopathology were studied in cases with pulmonary atresia and PDC [2].

The (histo)pathology of the human DA to left pulmonary connection of severe PDC is shown in neonatal post-mortem specimen with pulmonary atresia and tetralogy of Fallot. Clinical images of three human patients are presented in addition.

The use of the historic human cardiac pathology and embryo collection of the department of Anatomy and Embryology Leiden University Medical Center was approved by the local Ethics committee.

Informed consent was obtained from all patients or their parents.

### Immunohistochemistry

For histological examination, mouse embryos were processed [14, 22]. In brief, they were fixed in 4% paraformaldehyde (0.1 M, pH 7.4, 24 h, 4°C), embedded in paraffin, and serial sections (5 μm) were mounted on glass slides.

Deparaffinization was followed by ethanol and rehydration. Endogenous peroxidase activity was inhibited and slides were microwaved for antigen retrieval. Sections were incubated with primary antibodies against eGFP (Abcam ab 13970, 1/500), Tropomyosin (Sigma-Aldrich T9283, 1/500), TFAP2α (GeneTex GTX62588, 1/2000), NKX2.5 (Santa Cruz SC-8697, 1/4000), MEF2c (Cell Signaling Technology CST5030), αSMA (A2547 (Sigma-Aldrich, Missouri, USA) and PECAM1 (Santa Cruz SC-1506-R). Diluted primary antibodies (PBS–Tween-20 (PBST), 1% bovine serum albumin; A8022; Sigma-Aldrich, St Louis, MO, USA) prevented non-specific binding. Slides were rinsed twice in PBS and once in PBST. Tiramide signal amplification (TSA PLUS Biotin kit, NEL749A001KT, Perkin Elmer, Waltham, MA, USA) was used in NKX2.5 staining by adding HRP-labelled antibodies followed by amplification according to the manufacturers’ manual. For fluorescent microscopy primary antibodies were visualized by incubating in fluorescently labelled secondary antibodies (in PBST, 60 min). Secondary antibodies were diluted 1/200 and applied [14] DAPI (Life Technologies D3571, 1/1000) was used as nuclear stain while slides were mounted with Prolong Gold (Life Technologies).

The human material was stained for standard histology (Hematoxylin-eosin, Azan, Resorcin-fuchsin), furthermore, embryonic human material was stained immunohistochemically for HNK1 and HHF35 [25] and NKX2.5 (SHF) and TFAP2α (NCCs) [27].

### Microscopic analyses and 3D reconstructions

3D reconstructions of the arterial poles were made using Amira 6.3 (Template Graphics Software Inc., Houston, TX, USA) of mouse embryos between E10.5 and E16.5. Slides were scanned using the Pannoramic 250 Flash III slide scanner (3DHISTECH Ltd, Budapest, Hungary) and images of identical scale and exposure were exported using Histech Pannoramic Viewer (3DHISTECH Ltd.). Images were stacked and semi-automatically aligned in Amira. Relevant cardiac and arterial structures were labelled based on staining patterns and morphology. Surface views were exported to PDF formats using the Adobe Acrobat 9.5 software package.

## Results

### Normal development in the WntCre reporter mouse embryo

Embryos from E10.0 to E12.5 show the most important development and remodelling with a gradual change in the position and connections of the 4^th^ and 6^th^ PAA to both the aortic sac, later on separated into the ascending aorta and PT, and the dorsal aortae.

#### Pharyngeal arch artery and pulmonary artery remodelling

At E10.0 the 3^rd^ PAA, encased within WntCre positive NCCs, is connected to the aortic sac. Directly caudal of the 3^rd^ PAA, ventral sprouts of the 4^th^ PAA, covered by green labelled NCCs can be detected having a course more parallel to the dorsal aortae instead of the more transverse position of the 3^rd^ arch (Figure 2a,c,d). Small indications of dorsal sprouts, covered by green labelled NCCs (Figure 2c), are seen but no continuous 4^th^ or 6^th^ PAA is formed yet (Figure 2a,c,d). Mid-sagittally between these developing ventral sprouts of the PAAs there is an endothelial network, the MPES. Cranially, the endothelial network is surrounded by yellow labelled Nkx2.5 positive SHF while the more caudal extensions of the MPES are surrounded by non-Nkx2.5 stained pharyngeal mesenchyme (Figure 2b,d). Connections of the MPES to both the left and right ventral sprouts of the 4-6^th^ PAA can be discerned (Figure 2a,d).

**Figure 2:**
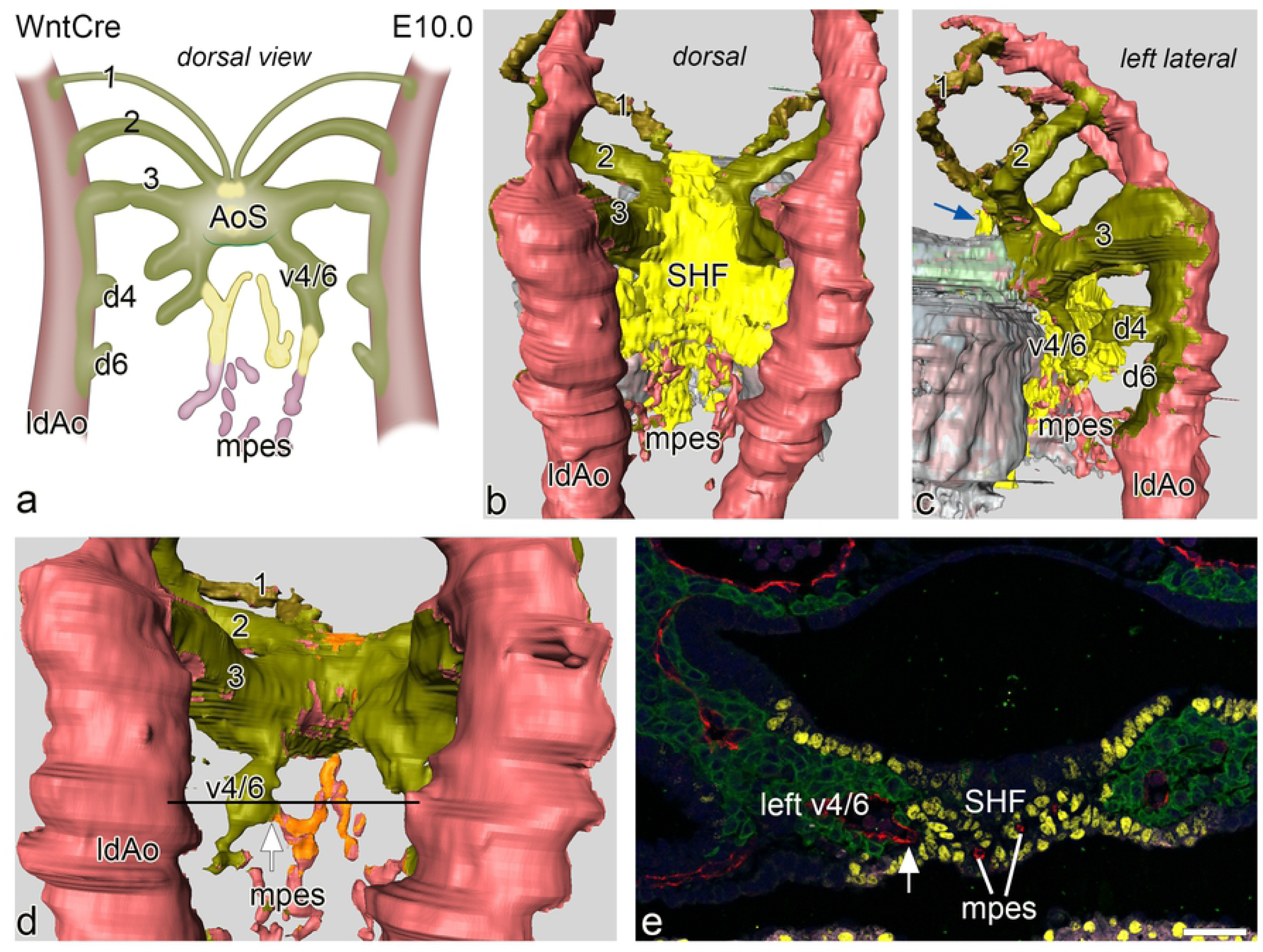
**a.** Schematic representation of a dorsal view of the connection of the vasculature of the mid-pharyngeal endothelial strand (mpes) and connections to the combined ventral sprouts (v4/6) of the right and left 4^th^ and 6^th^ PAA continuous with the aortic sac (AoS) in an E10.0 WntCre mouse embryo. The dorsal sprouts of 4^th^ and 6^th^ PAA (d4 and d6) are connected to the dorsal aorta (indicated is the left one, ldAo). Fig **b, c and d:** Reconstructions from various angles of relevant structures showing the WntCre positive neural crest cells (olive green) surrounding the continuous PAA arteries (1,2,3) as well as the AoS (out of view in **b.**) The dorsal aortae are on their lateral and dorsal aspect surrounded by NCC. The myocardium is depicted in shades of grey. The mid-line mesenchymal second heart field mass (SHF: yellow) is positioned between the confluence of the 3^rd^ (running anteriorly over the AoS; blue arrow in **c**) and the developing 4^th^ and 6^th^ PAAs (in **b, c**). Fig **d.** In this reconstruction the mid-line SHF mesenchymal mass has been removed showing the extent of coverage of the cranial part of the mpes (orange) while more caudally the mpes is not surrounded by SHF (red). Fig **e.** transverse section at the level (see black line in **d**) of the connection (white arrow) of the cranial part of the mpes (white lines) and endothelial cells (stained red for CD31) to the left v4/6 (NCC: green). The NKx2.5 positive SHF cells (yellow) border this connection and surround the endothelial cells here in the midline. Magnification: bar 100 µm.

At E11.5 the situation is changed in that the 4^th^ PAAs are now completely developed and connect the aortic sac with the dorsal aortae. Also the 6^th^ PAAs are complete (Figure 3a-d). The dorsal sprouts of the 6^th^ PAA are solely surrounded by NCCs and devoid of SHF (Figure 3a-e). At this stage the proximal parts of the future left and right pulmonary artery, that at E10.0 are covered by SHF, have disappeared and seem to be incorporated in the short ventral sprouts of the 6^th^ PAA. These are special in that they have a double lining with NCCs of the original 6^th^ PAA ventral sprout laterally, and Nkx2.5 SHF medially (Figure 3d,e) being continuous with the mid-line SHF mass (Figure 3b). The distal part of the pulmonary arteries is not surrounded by either SHF or NCC (Figure 3 a,b,d), but are embedded in Mef2c positive splanchnic mesoderm as shown in a E12.5 embryo (Figure 4k).

**Figure 3:**
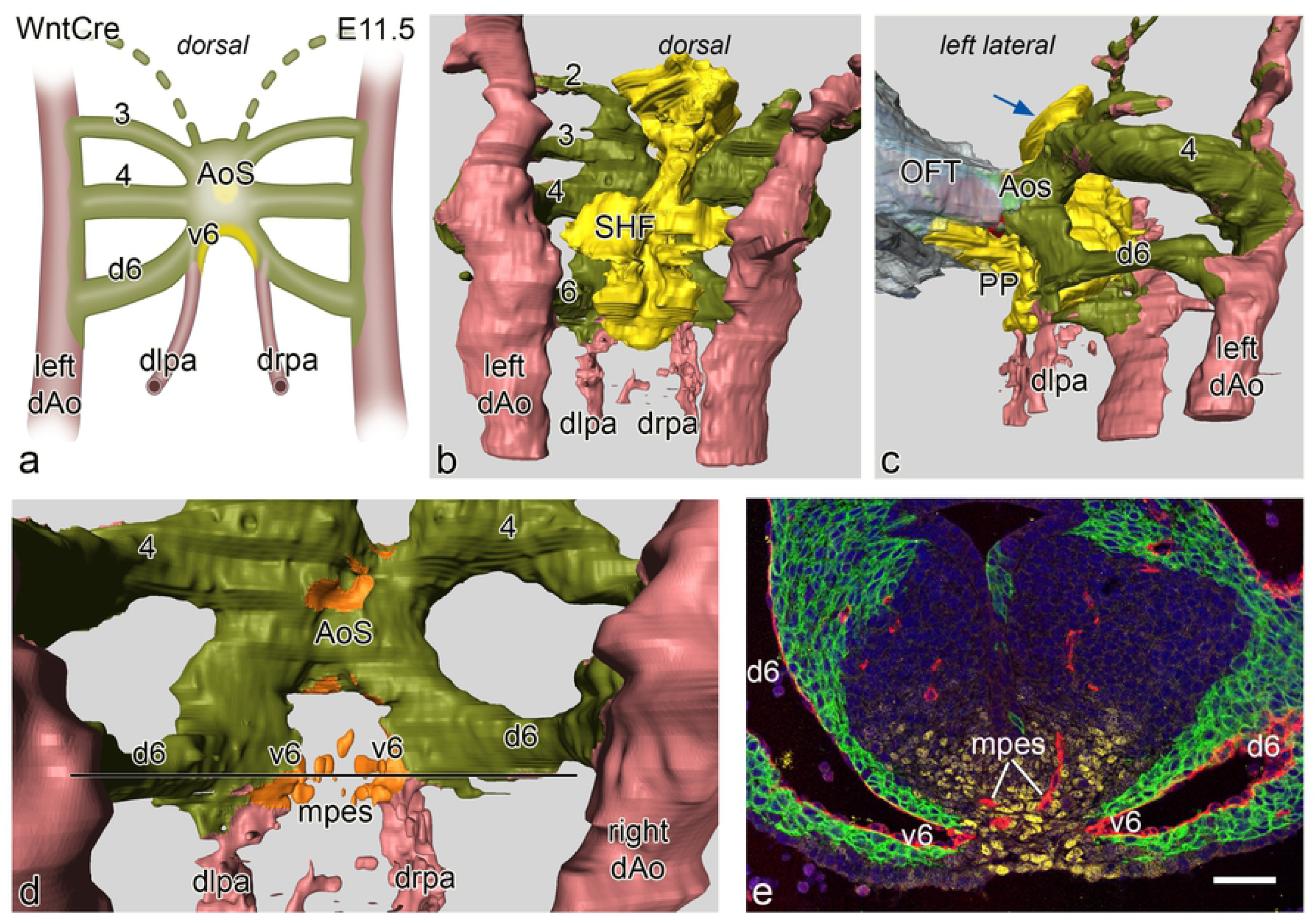
**a**. Schematic representation of a dorsal view of vascular connections in an E11.5 WntCre mouse embryo. In this stage PAA 3, 4 and 6 are complete. The distal left (dlpa) and right (drpa) pulmonary arteries merge sideways with PAA 6 creating a ventral (v6) and a dorsal (d6) segment of the latter. The d6 is completely encircled by neural crest cells (green) while the v6 has a lateral layer of neural crest and a medial layer of second heart field (SHF: yellow). The v6 forms a short proximal pulmonary artery. Fig **b**. dorsal view of a reconstruction at this stage with now complete 3^rd^,4^th^ and 6^th^ arches that are lined by neural crest (olive green). The SHF (yellow) forms the mid-line mesenchyme. The dlpa and drpa (pink) are not covered by either NCC or SHF. Fig **c.** left lateral view showing the connection of the outflow tract (OFT) myocardium (grey) with the short aortic sac (AoS). The SHF mass has a short extension anteriorly towards the future aortic orifice (blue arrow). More prominent at this stage is the left sided SHF extension that runs underneath the v6 and along the future pulmonary wall of the AoS. This is the so-called pulmonary push (PP). Fig **d.** dorsal view after removal of the SHF, the double lining of the relatively short v6 (identical to a proximal part of a pulmonary artery (see also **a.**) is depicted with a NCC (olive green) and a SHF reflected coverage (orange) Fig **e.** section of the embryo at the level indicated (black line) in **d**. The right and left v6 (endothelial cells red) are in part lined by NCCs (green) and Nkx2.5 positive SHF (yellow). In the SHF midline mass (yellow) the endothelial plexus (red) of the mid-pharyngeal endothelial strand (mpes) is visible. Magnification: bar 100 µm.

**Figure 4:**
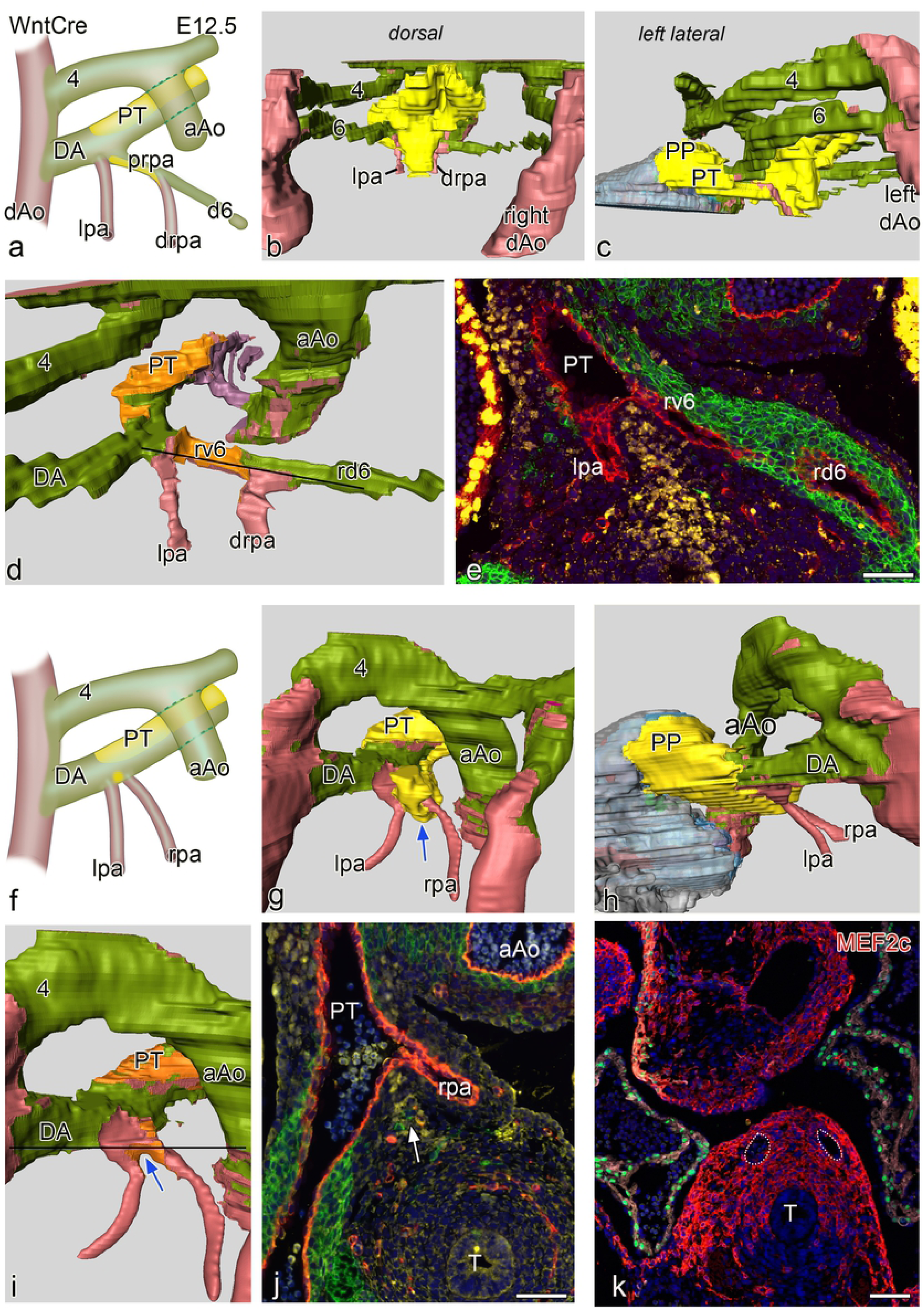
**a** and **f** schematic representations of two E 12.5 WntCre embryos. Embryo **a.** is slightly less far developed as compared to **f.** In both embryos the ascending aorta (aAo) and the pulmonary trunk (PT) have now been separated. Fig b-d shows reconstructions of the younger embryo with a complete 4^th^ and 6^th^ PAA. Fig **c** shows the dorsal midline SHF mass (yellow) partly covering the neural crest (green) lined PAA’s. Fig **c** is a left-sided view showing the position of the SHF mass (yellow) which has an extension (pulmonary push population: PP) covering the major part of the lumen of the PT towards the myocardial outflow tract (grey), running underneath the 6^th^ PAA to pulmonary artery connection. Fig **d**. dorsal view after removal of the SHF mass (lumen coverage area : orange). On the left side the dorsal 6^th^ PAA, now referred to as ductus arteriosus (DA) is completely surrounded by neural crest cells, There is no indication of a ventral 6^th^ PAA. The left pulmonary artery (lpa: red) abuts independently on the PT On the right side the situation is less well developed. The dorsal segment of the right 6^th^ PAA (rd6) is regressing and completely surrounded by NCC. The ventral segment of the 6^th^ PAA (rv6) is still present with a lateral wall of NCCs and a medial wall of SHF. The distal part of the right pulmonary artery (drpa) enters side-ways into this right-sided 6^th^ PAA. Thus the proximal rpa is at this stage formed by the rv6. Fig **e**. section at the level indicated in **d** (black line) in which it can be seen that the right v6 (rv6: also proximal part of the rpa) has both a lining of SHF (yellow) and NCCs (green). Fig **f** in the more developed embryo the originally distal parts of the lpa and rpa, embedded in Mef2c positive mesoderm (Fig **k**) are not covered by NCC and SHF. They enter the PT independent of the DA. Fig **g-i** are reconstructions showing similar dorsal views as **b-d.** Level of section **j**. is indicated by a black line in **i.** The SHF derived flow divider is still seen between the lpa and rpa (blue arrows in **g** and **i** and white arrow in **j**). The right d6 has regressed completely. There is on both sides no indication anymore of the v6 segments. **k** section of a E12.5 Mef2cCre embryo showing that the distal parts of the lpa and rpa (white dotted vessels) are situated within Mef2c positive splanchnic mesoderm that is not stained by Nkx2.5 (green). T: trachea Magnification sections **e,j,k** bars100 µm.

At E12.5 we present two embryos with a slightly different stage of development (Figure 4a-j). In both cases there is an OFT separation of the ascending aorta and the PT. The younger embryo is comparable to the E11.5 case with regard to development on the right side where, next to a regressing right dorsal 6^th^ PAA, the ventral 6^th^ PAA segment show a combined wall of NCC and SHF (Figure 4 a,d,e). The distal part of the right pulmonary artery, therefore, connect to the 6^th^ PAA between its dorsal and ventral segment (Figure 4d). This ventral segment of the 6^th^ PAA present as the proximal part of the right pulmonary artery. On the left side we can not discern a separate ventral 6^th^ PAA segment anymore as now the distal part of the left pulmonary artery, not being encased by NCC and SHF, enters directly into the PT. The left 6^th^ PAA, also referred to as DA, consists of the dorsal segment being solely surrounded by NCC (Figure 4 a,d). The mid-line SHF mass show a marked pulmonary push population along the PT.

In the older embryo (figure 4f-j) the ventral segments of the 6^th^ PAA (in the previous stages the proximal pulmonary arteries) are almost completely incorporated in the dorsal wall of the PT. The pulmonary arteries, however, not surrounded by either NCCs or SHF and embedded in Mef2c positive mesoderm (Figure 4 k), arise separately from the PT and even seem to form part of the dorsal wall of the PT (Figure 4 f,g,i). The right dorsal segment of the 6^th^ PAA has regressed completely while the dorsal left 6^th^ PAA persists as the definitive DA. The latter, demarcated because being surrounded by NCCs, is now positioned more anteriorly in continuity with the PT. The mid-line SHF mass has now dissolved and is mainly recognisable in the flow divider between the left and right pulmonary arteries as well as its extension as pulmonary push along the PT (Figure 4 g-j).

#### Role of the midline Nkx2.5 positive SHF population and the pulmonary push

The mid-line pre-pharyngeal Nkx2.5 positive SHF population, contacting the dorsal wall of the aortic sac, deserves additional attention. At E10.0 these cells extend anteriorly over the 2^nd^ and 3^rd^ PAA towards the aortic side of the outflow tract up to the borderline with the myocardium, reaching the site where the intercalated aortic valve leaflet will develop (Figure 2c). Between E11.5 and E12.5 a clear asymmetry develops in the Nkx2.5 positive SHF population. This becomes more prominent initially dorsally and thereafter along the left (pulmonary) side of the aortic sac (so-called pulmonary push). At E12.5, after separation of the OFT, the resulting population runs along the left side of the PT and reaches the myocardium at the site of the future pulmonary intercalated valve leaflet area (Figures 3 c, 4 c,g,h). This left-sided pulmonary push population runs below the NCC-surrounded DA. Exactly in the corner of this connection of DA with PT the left pulmonary artery enters the PT at the border of SHF and NCCs (Figure 4c,g,h). At the right side, where no pulmonary push SHF is present, the dorsal 6^th^ PAA segment (comparable to the left DA) regresses.

As a consequence the left dorsal segment of the 6^th^ PAA forms the DA while the right dorsal segment, which is already separated from the right pulmonary artery, regresses. Therefore, during normal development there remains after E12.5 no marked length of a ventral segment with a sideway connection of the pulmonary arteries.

### Normal development in human embryonic stages

A renewed study of human embryonic stages 4-24 mm CR length (n= 5, Carnegie stages: CS12-24) confirms the data from the mouse studies showing a similar remodelling and incorporation process of the short ventral sprout of the 6^th^ PAA and the attached left and right pulmonary arteries into the PT. The youngest embryo (4 mm, CS 12 of the Viennese collection) shows the connection of the MPES network to the developing combined 4^th^ and 6^th^ PAA ventral sprouts (Figure 5a). Somewhat more advanced configurations are shown in a 6.5 mm embryo (CS 14, Leiden collection) and a 9.5 mm (CS 16). In these embryos we observed only the morphologic configuration of the vessel lumen as no immunohistochemistry was available at the time of collection of this rare material. We now show, however, that in both embryos the distal pulmonary artery connects sideways to the 6^th^ PAA with a very short ventral 6^th^ PAA segment (Figure 5 b,c) comparable with normal mouse embryos. Access to more recently collected valuable embryonic material (CS 16) (Amsterdam UMC), in which immunohistochemistry was performed, confirms the presence of the Nkx2.5 pulmonary push population as well as the adjoining, non-overlapping TFAP2α positive NCC cells. The resolution and number of the sections does not allow for detailed reconstruction of this area or decisions on the presence of a ventral 6^th^ PAA segment (Figure 5d,e).

**Figure 5.**
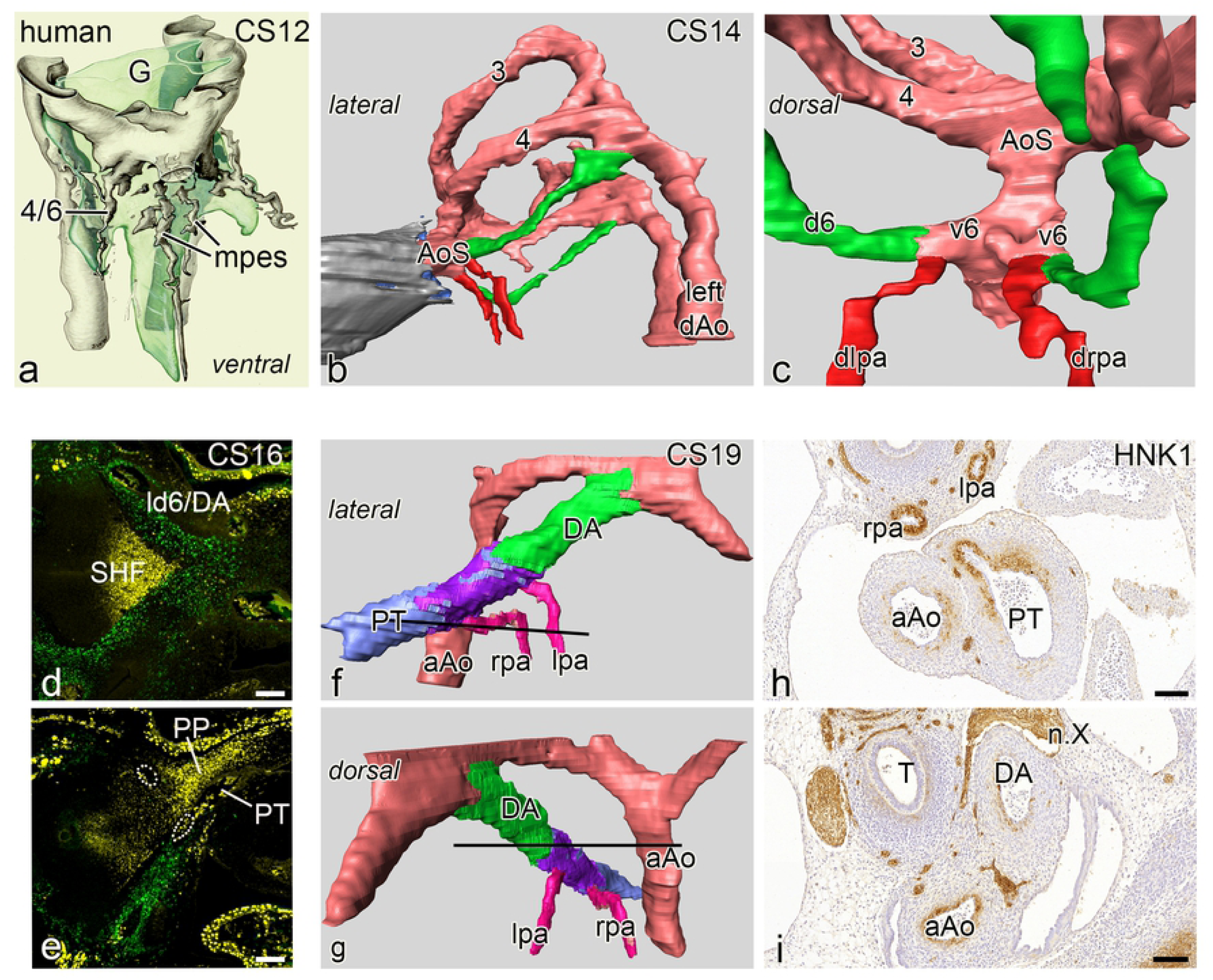
Human embryos in an upclimbing age series, Reconstructions of embryos depicted in **a,b,c** have been stained with hematoxylin eosin. **a.** Reconstruction of a very early human embryo (4 mm, CS 12) with a complete 3^rd^ PAA. The combined ventral sprouts of the 4^th^ and 6^th^ PAA are in contact with the mid-pharyngeal endothelial plexus (mpes) all depicted in grey and surrounding the developing gut (light green, G). Fig **b,c** left lateral and dorsal view of a lumen reconstruction of a human embryo (6.5 mm, CS14) with complete 3^rd^ and 4^th^ arches (3,4). The dorsal 6^th^ artery (d6) is indicated in green, the myocardium of the outflow tract in grey and the distal pulmonary arteries in red. The dorsal view the short ventral segments of the 6^th^ PAAs (v6) form a short proximal pulmonary artery. The distal part of the pulmonary arteries (dlpa,drpa:red) are thus in contact with the AoS. Fig **d,e.** sections of a human embryo (CS16) which is stained for neural crest for TFAP2α (green) which clearly surrounds the ld6/DA (**d**). Fig.**e.**There is a marked extension of Nkx2.5 positive SHF (yellow) along the outer side of the PT (pulmonary push population: PP). It cannot be discerned whether the lpa and rpa (white dotted rings) are of proximal or distal order. Fig **f-g** reconstruction of an older human embryo (17mm, CS19) stained for HNK1. The ascending aorta (aAo) and the PT have been separated. The right 6^th^ PAA has disappeared. The lpa and rpa (pink) are connected separately to the dorsal side of the PT, sharing the HNK1 antibody staining (purple) that allows for distinguishing boundaries between the left sided dorsal 6^th^ PAA (ductus arteriosus: DA, green) and the PT (light blue). The DA continues smoothly into the PT with a more anterior position as compared to the lpa and rpa. Fig **h,i**. transverse sections (see section levels as black lines in **f** and **g**) of the HNK1 stained pulmonary arteries (lpa and rpa) and the connection of the rpa to the PT, while the main part of the PT and the DA are negative for this staining. Abbreviations: n.X vagal nerve, T trachea. Magnification: bars 100 µm.

Study of an older embryo (Leiden collection, CS 19, 17mm), in which the boundaries of the pulmonary arteries, PT and the DA were visualized by immunohistochemical staining with an anti-HNK1 antibody, reveals a next stage of development. The differential staining of various SMC types is not primarily related to either NCC or SHF but shows a specific patterning at this stage. The pulmonary arteries are separately connected dorsally to the PT and in continuation with a ring-like part of HNK1 positive SMC between the DA and the PT (Figure 5 f-i). This area might be indicative of the incorporated ventral segment of the 6^th^ arch (Figure 5f,h). The dorsal sprout of the left 6^th^ PAA (referred to as DA), hardly stained for HNK1 (Figure 5 i), is more anteriorly in smooth continuity with the, also non-stained main part of the PT (Figure 5 h). The right 6^th^ PAA has already disappeared thus not encumbering the connection of the right pulmonary artery to the PT (Figure 5f,g).

### Pulmonary ductal coarctation (PDC) and interruption of the proximal left pulmonary artery

#### Clinical examples

In a case of pulmonary stenosis without VSD an angiography frame shows the normal connection of the leftsided DA above and anterior of the left pulmonary artery. The bifurcation of the the left and right pulmonary arteries, presenting no indications of narrowing at their origin, (Figure 6a) is more posterior. In two clinical cases with PDC there is an abnormal lateral DA to left pulmonary artery (Figure 6 c). In the first case, a neonate with tetralogy of Fallot with pulmonary atresia and confirmed 22q11 deletion, PDC is demonstrated by angiography (Figure 6 c). The second case with no detected genetic abnormality, shown by echo Doppler (figure 6h-k), concerns a DORV (tetralogy of Fallot type) with pulmonary atresia in whom the development of the severe obstruction of the proximal left pulmonary artery can be observed during the clinical course. The child was born at 31 weeks gestation with a weight of 1075 grams. With a widely patent DA on prostaglandin the left pulmonary artery origin is slightly smaller than the remainder of the branch (Figure 6h,i). At term (weight 2100g) a 3,5 mm modified Blalock-Taussig-shunt was connected to the right pulmonary artery and prostaglandin was discontinued. After DA closure the origin of the left pulmonary artery was nearly obliterated (Figure 6 j,k) and the child needed a second shunt to supply the left pulmonary artery. After uneventful recovery the child is now awaiting corrective surgery.

**Figure 6a.**
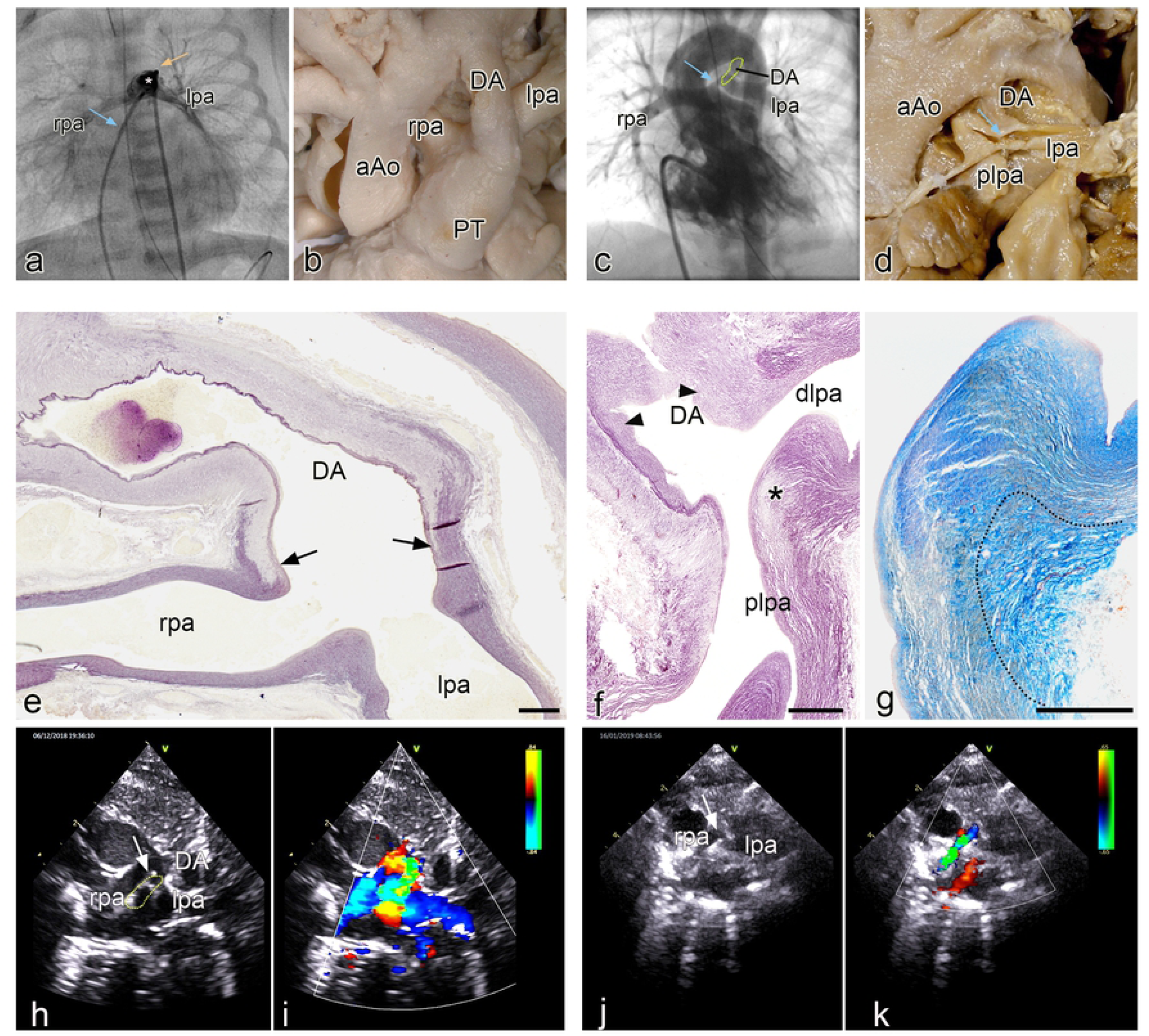
Angio of a patient with pulmonary atresia without VSD in which the leftsided DA (asterisk) is positioned above and anterior of the left pulmonary artery. The left (lpa) and right (rpa) pulmonary arteries (venous line :blue arrow; arterial line yellow arrow) are not compromised at their origin. Fig **b**. Morphology of the arterial pole of a heart specimen in which the ductus arteriosus (DA) connects anteriorly to the pulmonary trunk (PT), while the rpa and lpa are more dorsally connected. Fig. **c**. Angio of a patient with tetralogy of Fallot with a marked narrowing (blue arrow) of the proximal (origin) left pulmonary artery (DA dotted area). Fig **d** Postmortem specimen with atresia of the PT and a confluence of the rpa and lpa, with a smaller diameter of the latter. The DA enters sideways into the lpa which shows a PDC at the site of connection (blue arrow). Fig **e.** sagittal section of a human fetal (resorcine fuchsine stained for elastin). The elastin poor muscular DA only lined on the lumen side by an internal elastic lamina, connects with a fishtail like construction (arrows) to the elastin rich rpa and lpa. No DA tissue is encountered in the wall of the lpa and rpa. Fig.**f**. Sagittal section (resorcine fuchsin stained for elastin) of a specimen with severe pulmonary ductal coarctation. The tissue of the elastin poor DA, presenting with intimal cushions (arrowheads), is inserted sideways in the wall of the plpa (asterisk), while the dlpa is still elastic in nature. Fig.**g**. detail of **f** in a subsequent section stained for Azan in which it can be seen that the elastic lamellar structure (yellowish) of the dlpa and plpa is interrupted by the adventitia (dotted area) of the DA. **Fig h.** 2D echocardiographic image of the patient with DORV on prostaglandin. In this high parasternal short axis view the rpa, lpa and DA (dotted area) are indicated. The arrow points to the lpa origin. Fig **i.** Same image with Doppler color showing the flow to the rpa and lpa in blue and the DA flow in orange and green. **Fig j.** 2D echo of the same patient 10 days later after placement of the right mBT shunt and discontinuation of prostaglandin. Same view as Fig **h**. The DA has closed. The origin of the lpa is severely stenosed marked by the arrow. **Fig k.** same image as **i** with showing the flow to the rpa. Because of the mBTS the rpa the flow is turbulent coded in green. The lpa does not receive flow because of the severe stenosis at its origin. The DA has closed. Magnification: e-g bars: 100 µm.

#### Pathology in human neonates with PDC

The macroscopy and histopathology were investigated in 7 selected cases (Leiden congenital heart specimen collection) with a PDC (Figure 6 d,f,g) and compared with specimen with a normal morphology (Figure 6 b,e). The pathology specimen have been previously described [2] and were restudied for details of DA, PT and left pulmonary artery connection. All specimens had a pulmonary atresia (Figure 6 d) and in 6/7 also a VSD. In all cases the DA inserted sideways into the left pulmonary artery, creating a proximal (continuous with the PT) and a distal (running towards the lungs) part (Figure 6 d). The left pulmonary artery was often smaller than the right pulmonary artery (Figure 6 d). In all seven specimens the ductal tissue formed the major part of the obstructed proximal left pulmonary artery with a variable extension into this arterial segment. It was observed that the elastic lamellae of the media of the proximal and distal pulmonary artery were not continuous at the site of the PDC (Figure 6 f.g). In 2/7 there was a total occlusion of the left pulmonary artery by this ductal tissue. The lumen of the DA was in open continuity with the distal part of the left pulmonary artery, allowing for filling of the lungs. These histological findings were in marked contrast to the normal neonatal specimen where the elastin-poor DA connected, with a fish tail like construction to the elastic PT and the adjoining left and right pulmonary arteries (Figure 6 e).

### Abnormal development of the pulmonary artery, DA and PT connection in the VEGF 120/120 mutant mouse

To obtain more insight in mechanisms underlying aberrant DA to pulmonary artery connections we studied a series of VEGF 120/120 mutant mouse embryos and their development-matched wild type embryos. New in this paper compared to the hitherto described VEGF series [22, 23] was the addition of mouse embryos in which we could discern the SHF population by staining for Nkx2.5 and thus the possibility to investigate in more detail the connection site of the PT, pulmonary arteries and 6^th^ PAA. The Nkx2.5 staining proved to be reliable at E10.5 and E11.5. Older stages from E12.5 onwards the staining was not conclusive anymore because of loss of staining intensity. So, we have focused on these earlier stages to understand more about the possible underlying mechanisms leading to the associated heart malformations.

Wild type mouse embryos at E10.5 (n:5, all stained for Nkx2.5) showed a marked presence of the dorsal mid-sagittal SHF population with a pulmonary push extension at the dorsal site of the aortic sac (the future PT side). Both 4^th^ PAAs were complete. The developing pulmonary arteries were connected to the ventral sprouts of the 6^th^ PAA which were not yet connected to the dorsal sprouts to form a complete 6^th^ PAA (Figure7a) and are embedded in this area in the Nkx2.5 positive SHF population.

At E11.5 the VEGF wild-type embryos were comparable to older E12.5 WntCre reporter embryos (Figure 4f-j), with a completion of the 6^th^ PAA and separation of the PT and the ascending aorta. The left 6^th^ PAA (now referred to as DA) was complete while the right 6th had already regressed.

In the VEGF120/120 E10.5 series (n=5; 3 stained for Nkx2.5) the major difference compared to the wild type, was found in the stunted appearance of the SHF population lacking a pulmonary push extension (Figure 7b). The connection of the pulmonary arteries was embedded in the SHF population and connected to the ventral sprouts of 6^th^ PAA like in the wild type embryo.

**Figure 7.**
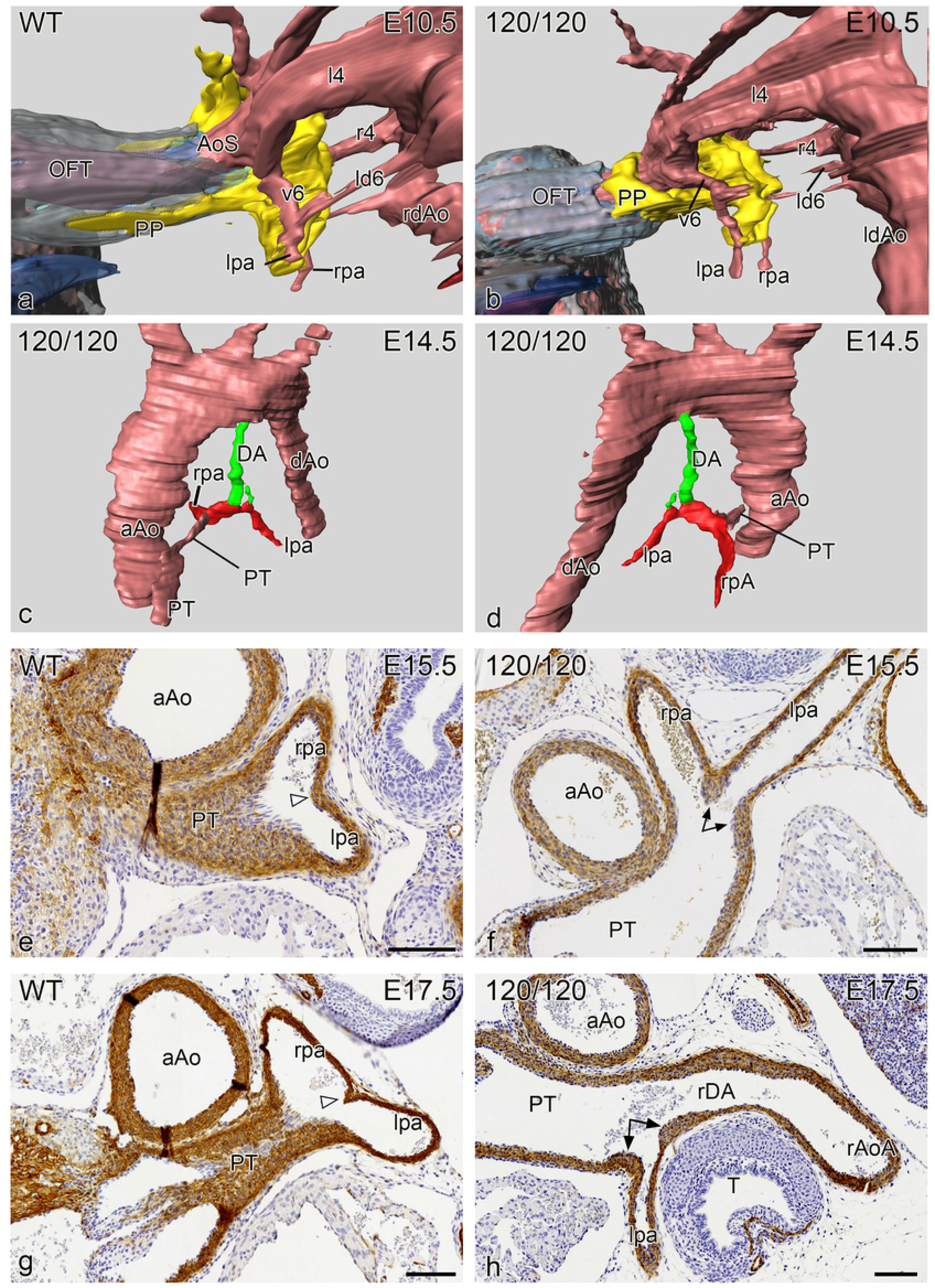
Reconstruction (left lateral views) of a E10.5 VEGF WT **(a)** and a VEGF 120/120 **(b)** mouse embryo in which the SHF has been stained for Nkx2.5 (yellow). There is a marked difference in the pulmonary push (PP) which extends along the future pulmonary side of the aortic sac (AoS) for quite some distance in the WT embryo (**a**), while in the mutant embryo (**b**) the PP is stunted. In both embryos the lpa and rpa connect to the AoS by way of ventral sprout of the 6^th^ PAA (v6) and are embedded in the SHF tissue**. Fig c,d** reconstructions of an E14.5 mutant embryo in which the hypoplastic DA (green), arising vertically from the aortic arch, is connected side-ways to the lpa (red). There is atresia of the pulmonary orifice and the pulmonary trunk (PT) is hypoplastic**. Fig e-h** Sections stained for α smooth muscle actin (1A4) of the site of connection of the lpa and rpa into the PT. **e** and **f**. are E15.5 embryos in which (**e**) the WT shows equal sizes of the lpa and rpa with a small posterior separating ridge (open arrowhead), while in the mutant embryo (**f**) the origin of the lpa, which is relative smaller in diameter as compared to the rpa, is flanked by thickened DA tissue (arrows). In a WT (**g**) and mutant embryo (**h**) of E 17.5 we observe a similar phenomenon although in the mutant embryo (**h**) the small lpa, encircled by thick ductal tissue (arrows) is connected to a right-sided DA (rDA) continuing into a right aortic arch (rAoA). Magnification bars in **e-h** 100 µm.

The VEGF120/120 mutant embryos at stage E11.5 (n=7; 3 stained for NKx2.5) all showed a complete 4^th^ and 6^th^ PAA as in the wild type. While at E10.5 the main differences were observed solely in the PP population, this situation had changed at E11.5 showing a variety of abnormalities. The SHF population was stunted and did not show a proper pulmonary push extension. The anlage of the left and right pulmonary arteries bordered the SHF population as in wild types. Both the right ventricular OFT and the PT were stenotic. With regard to the development of the 6^th^ PAA we observed a lack of proper left /right dominance. In 2/7 cases the right pulmonary artery merged sideways with the right 6^th^ PAA, which thus still showed a ventral and a dorsal part. In one case the right dorsal 6^th^ PAA formed a right-sided DA continuing in a right aortic arch. In all cases the left pulmonary artery connected already separately to the PT while the left dorsal 6^th^ PAA (the regular left DA) was hypoplastic in 3/7 of these cases although, as yet, still in open connection with the hypoplastic PT.

At E12.5 the VEGF120 wild-type embryos (n=8) showed a comparable morphology to E11.5. The VEGF 120/120 embryos (n:5) showed similar abnormalities as described for E11.5, complemented by a case with a retro-esophageal left subclavian artery and a case with a sling configuration of the arch.

In the VEGF embryos of E14.5 – E18.5 (n=15) we specifically looked for cases with an incomplete incorporation of the ventral sprout of the 6^th^ PAA which leads to an abnormal left pulmonary artery-oriented insertion of the DA seen in human patients with PDC (Figure 6 e). In one case this was very markedly combined with an hypoplastic left DA (Figure 7c,d). In four other cases we found thickened DA tissue encircling the entrance of the left pulmonary artery into the PT (Figure 7e-h).

## Discussion

PDC is relatively common in cases with right ventricular outflow tract stenosis and cyanosis including tetralogy of Fallot with a severe stenosis to a completely atretic pulmonary orifice and trunk [2, 3, 28, 29, 30]. These anomalies are often seen in combination with the 22q11 deletion syndrome [31, 32, 33]. PDC has been postulated to be the counterpart of the aortic coarctation [29] in which DA tissue encircles the inner part of the aortic arch for more than 50%. However, the elastic structure of the aortic wall is always continuous as an outer layer from the aortic arch to the descending aorta [34]. The exact background of the development of a preductal aortic coarctation is still lacking. However a hemodynamic component has regularly been postulated [29, 35, 36] related to an augmented flow through the DA during development of the stenosis to atresia of the left ventricular outflow tract or aorta [28, 29, 36]. Waldman et al [20] studied patients with developing proximal left pulmonary artery interruption including the histopathology of the stenotic/interrupted area. They showed that in PDC the DA tissue could form the complete wall between the proximal and distal part of the left pulmonary artery. Reassessment of the histopathology in our initially described material [2] showed this to be correct. At the time we mistook the adventitia of the DA for a continuous outer media and had erroneously postulated a hemodynamic cause, comparable to aortic coarctation. We now realize that the connection of the DA to the proximal part of the left pulmonary artery is a developmental anomaly, as a in general the DA never encroaches on a pulmonary artery but connects separately to the PT forming a continuous arch towards the aorta as we showed in this study for the later stages in both normal human and mouse development.

The above background and findings shed a new light on the development and morphology of the interrelationship of the (left) pulmonary artery, the PT and the DA.

First, the investigations in subsequent series of early human embryos and comparable rat and chicken stages refined earlier interpretations. The 6^th^ PAA indeed develops from a ventral sprout from the aortic sac, connecting to the pulmonary artery vasculature, not by sprouting towards the lung parenchyme [7] but by recruiting endothelial cells from the MPES [10]. We demonstrated that mouse and human embryos were comparable and that both showed a short period in which there was a ventral and dorsal segment of the 6^th^ PAA. Only in older stages the pulmonary arteries connected directly to pulmonary trunk. The option that the pulmonary arteries directly connect to the aortic sac, as postulated [37] are a matter of semantics as in their definition the ventral segments appear to belong to the aortic sac.

Second, currently available (transgenic) reporter mouse data, as used in this study, allowed us to investigate in detail the contribution of NCCs and SHF cells (Nkx2.5 positive) at the hub of the various involved vessels. The ventral segments of the 6^th^ PAA, which have a dual lining of NCC and SHF, become incorporated in the posterior wall of the pulmonary trunk, after which the definitive pulmonary arteries, embedded in Mef2c positive mesoderm, have an independent connection to the posterior wall of the pulmonary trunk. The right and left dorsal 6^th^ PAA segments are completely surrounded by NCCs as has been shown before [12]. Normal timing of events allows the right 6^th^ PAA to regress, not narrowing the orifice of the right pulmonary artery to PT connection, while the left 6^th^ PAA becomes the DA with a smooth anterior connection to the PT. These data support the recent observation that the mesoderm surrounding the definitive pulmonary arteries lack Tbx1and thus Nkx2.5 SHF expression [38], explaining the absence of a direct effect of the 22q11 deletion on the development of the left and right pulmonary artery system.

The above findings reflect on the (histo)pathology in the human infant with proximal pulmonary artery stenosis. Defective formation of the pulmonary trunk, with absence of proper incorporation of the ventral 6^th^ PAA segment could lead to a situation in which the DA enters side-ways into a pulmonary artery.

Most patients with PDC and pulmonary atresia present with a VSD [30] indicating an early embryonic onset of the disease. In case of valvular pulmonary stenosis or atresia without VSD occurrence of PDC is rare [2] and the DA is positioned normally between the left and right pulmonary arteries. A number of patients with PDC present with a complete insertion of DA tissue between the proximal and distal part of the pulmonary artery. We considered a possible developmental mechanism, as a simple hemodynamic solution, with preferential retrograde flow from the aorta to the pulmonary trunk during developing stenosis to atresia of pulmonary trunk and orifice, proved not to be adequate. The available VEGF 120/120 mutant mouse model allowed us to study in more detail the DA-PT and pulmonary artery connection. It has already been reported [23] that originally both 6^th^ PAA were present and along with the developing PT stenosis to atresia, the left sided DA could also disappear. This was accompanied in some cases with development of additional aorto-pulmonary collateral arteries (MAPCAs). We now added that the most prominent abnormality was a stunted SHF population in early stages of mutant embryos. The PT did not receive the SHF cells as normally contributed by the pulmonary push population [18]. Thereafter, we observed lack of proper incorporation of the ventral segments of the 6^th^ PAA together with a loss of normal left-right dominance. This may result in a side-ways connection of a DA to the pulmonary artery. The loss of left-right dominance, with more frequent occurrence of a persisting right DA and a right aortic arch, as encountered in the mutant VEGF embryos, was also described in our human histopathology series [2] and also in patients with tetralogy of Fallot, pulmonary artery atresia and the 22q11 deletion syndrome [31, 32, 39]. However, the anomaly is not exclusive for this syndrome but seems to be linked to development of pulmonary stenosis to atresia in which it is encountered in 40% of the cyanotic infants with right ventricular outflow tract obstruction [3, 30]. In case of absent DA no PDC was seen to develop [30] as also reported in our (histo)pathology series [2] and confirmed in the VEGF model [23]. In both latter studies we observed a diminished diameter of the distal left pulmonary artery. In the VEGF embryos the hypoplasia of the left pulmonary artery occurred relatively late during development when the PT and the DA already showed hypoplastic characteristics. Thus, a secondary hemodynamic influence has been postulated [23]. A primary Tbx1 haploinsufficiency [40] was proposed for the diminished size of the left pulmonary artery in a study of the human 22q11 population, without additional cardiovascular anomalies, supported by data from Tbx1 heterozygous mice [38]. As, however, the definitive pulmonary artery (distal anlage) is not surrounded by Nkx2.5 positive cells (this study) nor by cells expressing Tbx1 [38], or Tbx1–dependent Hox genes [37] a secondary hemodynamic cause for this hypoplasia seems more obvious. After removing the obstruction in the proximal pulmonary artery there can be a catch up growth of the left pulmonary artery system [3].

For understanding the clinical presentation of PDC it is very relevant to know that neural crest-derived muscular DA tissue may continue contraction and stenosis formation even after birth. This process can be held responsible for postnatal development of proximal left pulmonary artery interruption. Surgical or (stent) interventions of PDC should ensure that DA tissue is not present anymore or can do no harm [3, 19, 20].

Since the majority of patients with PDC will be treated with prostaglandins (PGE1) to maintain DA patency before surgery the development of an interruption will be postponed till PGE1 is withdrawn after surgery. Although stenting of this segment may solve the problem initially it may in the long run be inferior to primary resection of all DA tissue.

## Acknowledgements

Funding and salary support of JP was provided by a grant of The Netherlands Heart Foundation (Projectcode: 31190BAV).

Dr.Aleksander Sizarov (Department of Pediatrics, Maastricht UMC, the Netherlands) and prof. Antoon Moorman (Department of Anatomy, Amsterdam UMC, The Netherlands) provided the human CS16 embryo.

The medical illustrator Ron Slagter is acknowledged for his excellent art-work.

